# CRISPR-Cas9 ribonucleoprotein-mediated co-editing and counterselection in the rice blast fungus

**DOI:** 10.1101/349134

**Authors:** Andrew J. Foster, Magdalena Martin-Urdiroz, Xia Yan, Sabrina Wright, Darren M. Soanes, Nicholas J. Talbot

## Abstract

The rice blast fungus *Magnaporthe oryzae* is the most serious pathogen of cultivated rice and a significant threat to global food security. To accelerate targeted mutation and specific gene editing in this species, we have developed a rapid plasmid-free CRISPR-Cas9-based gene editing method. It has previously been reported in *M. oryzae* that transformation with plasmids expressing Cas9 can generate specific mutations using sgRNAs, directing the endonuclease to specific genes. We show, however, that expression of Cas9 is highly toxic to *M. oryzae*, rendering this approach impractical. We demonstrate that using purified Cas9 pre-complexed to RNA guides to form ribonucleoproteins (RNPs), provides an alternative and very effective gene editing procedure. When used in combination with oligonucleotide or PCR-generated donor DNAs, generation of strains with specific base pair edits, in-locus gene replacements, or multiple gene edits, is very rapid and straightforward. Additionally, we report a novel counterselection strategy which allows creation of precisely edited fungal strains that contain no foreign DNA and are completely isogenic to the wild type. Together, these developments represent a scalable improvement in the precision and speed of genetic manipulation in *M. oryzae* and are likely to be broadly applicable to other fungal species.

---

In recent years, the use of the clustered regularly interspaced short palindromic repeats (CRISPR)-associated RNA-guided Cas9 endonuclease, has facilitated gene editing technologies have become the leading tool used to generate specific changes to DNA sequences in a wide range of species (1).

Within fungi, CRISPR-Cas9-based gene editing has been reported in many industrially relevant or model fungal species (2). The beauty of CRISPR systems lie in their simplicity: with most systems in current use possessing just two components to induce double stranded breaks (DSBs) in the genome of a target organism (3, 4). The first component is the Cas9 endonuclease, which cleaves target DNA at a genomic target sequence (5), while the specificity of the system is due to the second component, a single crRNA:tracrRNA chimeric guide RNA (gRNA), a single RNA molecule which in the CRISPR-Cas9 system uses a linker sequence to join the nuclease-binding tracrRNA and the target specific crRNA molecules found in naturally occurring complexes in the source organism *Streptococcus pyogenes* (6). The sgRNA associates with the nuclease and directs it to its genomic target sequence by sequence complementarity in the protospacer region, a short 17-20 bp sequence (6). The DSB created by the nuclease can then be repaired by non-homologous DNA-end joining (NHEJ) or using homologous recombination (HR), by introduction of donor DNA homologous to the sequence around the break, which allows very specific edits to the DNA sequence, or very precise insertions or deletions (7). The only target sequence requirement necessary for CRISPR-Cas9 gene editing is the presence of the protospacer adjacent motif (PAM), a triplet NGG located immediately 3’ of the genomic target sequence (8, 9). Because HR-based repair can be used to introduce modifications at some distance to the DSB, for example up to 30 bp in human stem cells (10), the majority of fungal genomes are accessible to manipulations using CRISPR-Cas9 editing.

CRISPR-Cas9 gene editing offers huge potential to accelerate the pace of research in key fungal research areas, such as biotechnology, medical mycology and plant pathology, by dramatically reducing the time required to undertake common objectives, such as targeted gene deletion, overexpression, or tagging the products of genes of interest with fluorescent proteins (2). CRISPR-Cas9 gene editing can also allow introduction of single nucleotide changes, facilitating rapid creation of multiple alleles for genes of interest. The technique also permits targeting of gene families, making multiple mutations (11) and studying dikaryotic, or polyploid fungi (12). The potential also exists to carry out ‘selectable marker-free’ manipulations for precise genetic changes, a prerequisite for any commercial application. CRISPR-Cas9 generated edible mushrooms have already, for instance, bypassed the gene manipulation regulations to which crop species engineered by methods preceding CRISPR were subject (13).

In the rice blast fungus *Magnaporthe oryzae*, a CRISPR-Cas9 gene editing system based on expression of Cas9 and CRISPR components *in vivo* has been reported to target single genes (14). However, the generation of mutants using this procedure has not been widely adopted and the protocol requires labour intensive cloning strategies so that the deletion of multiple genes would be not be practical. We therefore set out to look for an alternative method which might extend the range of applications for CRISPR-based gene editing technologies in this economically important pathogen of rice. Using purified nuclear-localised Cas9 (Cas9-NLS) and *in vitro* synthesised sgRNA, an approach pioneered in *Caenorhabditis elegans* (15) and subsequently in human cells (16) and fungi (17), we have been able to develop a ribonucleoprotein-CRISPR-Cas9 (RNP-CRISPR-Cas9) system. This procedure generates highly efficient rates of mutation in *M. oryzae* at a genomic target sequence, when a donor DNA carrying a selectable marker sequence and capable of repairing the DSB is co-transformed with the RNP into fungal protoplasts. Because we found that RNP-CRISPR-Cas9-mediated introduction of mutations was relatively inefficient without a donor DNA, we established what we term a gene co-editing strategy. This approach allows single nucleotide edits to be made without any other changes in or around a given target locus. Co-editing works by RNP-CRISPR-Cas9-mediated introduction of an oligonucleotide donor DNA, making a single nucleotide edit that confers resistance to an antifungal compound and simultaneous introduction of a second RNP and donor DNA that targets a second locus. In this way a useful proportion of antibiotic-resistant transformants can be identified that are edited at a second target locus. Additionally, a novel selection strategy that exploits negative cross resistance to two fungicides has been established to enable CRISPR mediated counterselection. This counter-selection method allows mutants to be created that are isogenic to an original wild-type strain. We believe that using RNP-CRISPR-Cas9 will permit precise and rapid gene manipulation in *M. oryzae* and other fungi, and thereby accelerate the pace of research in this economically important plant pathogen.

## Results

### Evidence of toxicity of Cas9 in *Magnaporthe oryzae*

Initially we reasoned that a *Magnaporthe* strain stably expressing a Cas9 gene would be a useful resource for the *Magnaporthe* research community, especially given the results reported by Arazoe and co-workers (14). We made multiple attempts to generate such a strain by introducing a codon-optimised Cas9 gene, together with a small guide RNA (sgRNA) targeting the melanin biosynthetic polyketide synthase-encoding gene *ALB1* (MGG_07219). Mutation of *ALB1* gives rise to an easily identifiable white (albino) colour phenotype in fungal colonies (18). We additionally made a separate construct to target a second melanin biosynthetic gene, *RSY1* (MGG_05059), which encodes scytalone dehydratase enzyme in which mutation gives rise to orange-red (rosy) fungal colonies (18). Despite many attempts, and using several different versions of Cas9 under control of different promoters, we were never able to generate mutants showing altered pigmentation, among the few transformants which resulted from transformations with either vector. Importantly, we were not able to reproduce the generation of mutants reported previously, even when the same vectors were used (14). We did, however, observe that the transformation of all constructs containing Cas9-encoding sequences always gave rise to far fewer transformants than empty vector controls (Fig. 1a and Table S1.). We conclude that stable expression of Cas9 is likely to be very toxic to *M. oryzae,* precluding widespread adoption of this method of gene editing.

### CRISPR-Cas9 gene editing using purified Cas9 and sgRNAs

In view of the toxicity of Cas9 nuclease to *M. oryzae* cells, we set out to assess whether CRISPR-Cas9 gene editing might instead be accomplished by introduction of purified Cas9 protein along with *in vitro* synthesised sgRNAs, into protoplasts of a wild type *M. oryzae* strain, Guy11. This approach has the advantage that the active CRISPR complex will only be transiently present in the fungus. To this end, we purchased nuclear-localised Cas9 (Cas9-NLS) from commercial suppliers (see Methods for details) and complexed this to gRNAs capable of directing Cas9 to the *ALB1* locus (Fig. 1b). We independently tested a second RNP that targets the *RSY1* locus. We introduced these Cas9-NLS-gRNA ribonucleoprotein complexes (RNPs) independently into a wild type *M. oryzae* strain Guy11,together with donor DNAs which would introduce an insertion containing a selectable marker (*HPH* − the hygromycin phosphotransferase gene cassette) near the 5’ end of the coding regions of these genes, by repair of the DSB by homologous recombination with donor DNAs containing at least 450 bp homologous regions on either side of the selectable marker (Fig. 1c). In both cases the selectable marker was expected to integrate close to the DSB−about 44 bp from the typical breakpoint in the genomic target sequence (the DSB is normally 3-4 bp from the PAM site) in both donor DNAs (Fig. 1c). We were able to demonstrate very efficient targeting of both genes as shown in Fig 1d and Table S2. Remarkably, mutation of *RSY1* was near to 100% efficient in multiple experiments and the efficiency of targeting *ALB1* was also greater than 50% in every test, with typically 70-80 % albino transformants generated (Fig. 1d and Table S2). As expected, the rates of mutation using donor-only controls were more typical of rates reported for gene deletion using conventional gene deletion strategies achieved by PEG-mediated transformation of protoplasts (see Table S2). These observations argue convincingly that RNP-CRISPR-Cas9 generated DSBs strongly induce HR repair and can therefore be exploited for efficient gene manipulation in *M. oryzae*.

**Figure 1.**
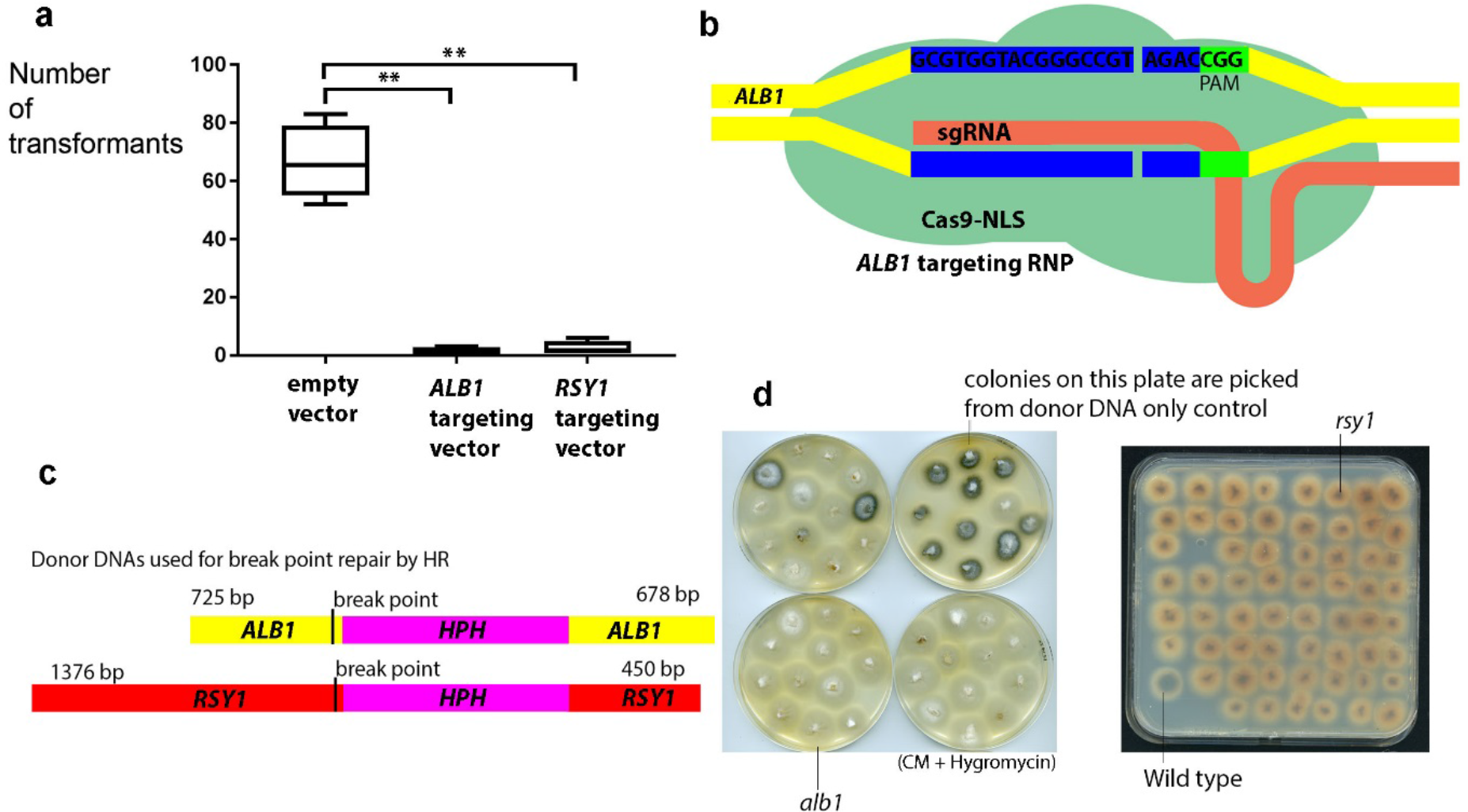
Toxicity of stably expressed Cas9 and efficacy of RNP-CRISPR-Cas9 gene targeting for the *ALB1* and *RSY1* genes. **a**. Binary vectors containing the gene encoding Cas9-NLS under the control of the *TrpC* promoter and terminator were introduced into Guy 11 using *Agrobacterium*-mediated transformation. Transformant numbers were assessed after 7 days on selective medium (transformants were subsequently sub-cultured for assessment of pigmentation after growth on CM). **b**. Illustration of the genomic target sequence for the *ALB1*-targeting RNP-CRISPR-Cas9 complex used showing a typical double stranded break in relation to the PAM site. **c**. An illustration of the donor DNA sequences used to repair DSBs created by RNP-CRISPR-Cas9 complexes and showing the position of break point relative to the selection marker hygromycin-phosphotransferase (HPH) used. **d**. Transformants picked from *ALB1*-targeting RNP-CRISPR-Cas9 + donor DNA transformation plates (before pigmentation was normally apparent) and growing on CM + hygromycin showing albino mutants and also (top right) transformants picked from a donor only control plate. On the right hand plate (square plate) are transformants picked from *RSY1*-targeting RNP-CRISPR-Cas9 + donor DNA transformation plates (before pigmentation was apparent) and growing on CM + hygromycin showing rosy (normally orange-red) pigmentation (one wild-type pigmented transformant is also present bottom left hand side of the plate).

### Highly efficient mutation of genes using CRISPR-Cas9 induced recombination of micro-homologous donor DNAs

The efficient mutation of *ALB1* by CRISPR-Cas9 induced homologous recombination of a donor DNA, prompted us to define the minimum length of homologous DNA that would facilitate efficient gene editing. Recent reports in some fungi suggest that micro-homologous regions are sufficient to allow repair by short donor DNAs of CRISPR-induced DSBs with high efficiency (19, 20). To test whether this was possible in *M. oryzae*, we amplified the *BAR* gene which confers resistance to the herbicide glufosinate ammonium (21), with 30 bp and 40 bp flanking regions on either side of the selectable marker (Fig S1). In this way we were able to demonstrate that a 30 bp region of homology was sufficient to induce repair by HR of the CRISPR-Cas9 generated DSB and result in mutation of *ALB1*, with efficiencies approaching those achieved with the much longer donor DNAs (see Table S3). The increased rates of mutation compared to those observed with donor-only controls made without RNP complexes, furthermore provided evidence that RNP-CRISPR-dependent gene replacement is efficient in *M. oryzae*. The use of such small flanking regions also demonstrates that RNP-CRISPR-Cas9 gene inactivation can be generated without the need for laborious cloning strategies.

### Direct demonstration of marker free mutation of the *ALB1* melanin biosynthesis gene using donor free CRISPR-Cas9 RNPs

Our observation of efficient gene replacement when RNPs were introduced into *M. oryzae* with donor DNA fragments, suggested that the Cas9-NLS enzyme creates DSBs and, in so doing, induces repair by the homologous recombination pathway. However, because these experiments used donor DNAs to repair the breaks using a selectable marker gene, they were genotypically indistinguishable from *alb1* or *rsy1* strains that would be generated from an experiment using the donor DNA only to disrupt each gene− as in ‘traditional’ gene disruption approaches. We therefore set out to examine CRISPR events more directly by introduction of the RNP complex targeting *ALB1* only, without donor DNAs to direct repair by homologous recombination. In the absence of donor DNA, the resultant DSB can be repaired by the non-homologous DNA end-joining (NHEJ) pathway which, because it is frequently inaccurate, should result in *alb1* (albino) mutants. Although such events were found to be rare under the conditions tested, we were able to observe albino mutants among a large background (showing confluent growth) of normally pigmented regenerants (Fig 2a). Control transformations without RNPs yielded no such white patches. Unfortunately, multiple attempts to dilute the transformants to a concentration where 10-20,000 individuals could be isolated, failed to yield any albino colonies, suggesting that much less than 0.01 % of regenerated protoplasts harbour *alb1* mutations. These observations suggest that the efficiency of the delivery system used would make marker-less gene targeting impracticable in the absence of an easily identifiable phenotype, because this would necessitate analysis of more than 20,000 individuals. Nevertheless, the albino mutants generated allowed us to directly demonstrate that RNPs are functional *in vivo* and to understand the nature of mutations generated through NHEJ. To this end, we purified albino mutants by several rounds of subculture followed by single spore isolation (Fig. 2a) and in a few cases, the insertion or deletion was large enough to be apparent by gel electrophoresis of the amplicons (Fig. 2b). In one case, no amplification was possible, indicating that a larger deletion had removed the sites where one or both primers, anneal. Sequencing of DNA around the genomic target site of the RNP in these albino mutants revealed true CRISPR mutations, showing a range of insertions or deletions close to the PAM site, as shown in Fig. 2c. These results demonstrated that the RNP complex functions by creating a DSB at the expected site and that marker-less single mutations are feasible, but not at a frequency which would be useful in the absence of an easily identifiable phenotype.

**Figure 2.**
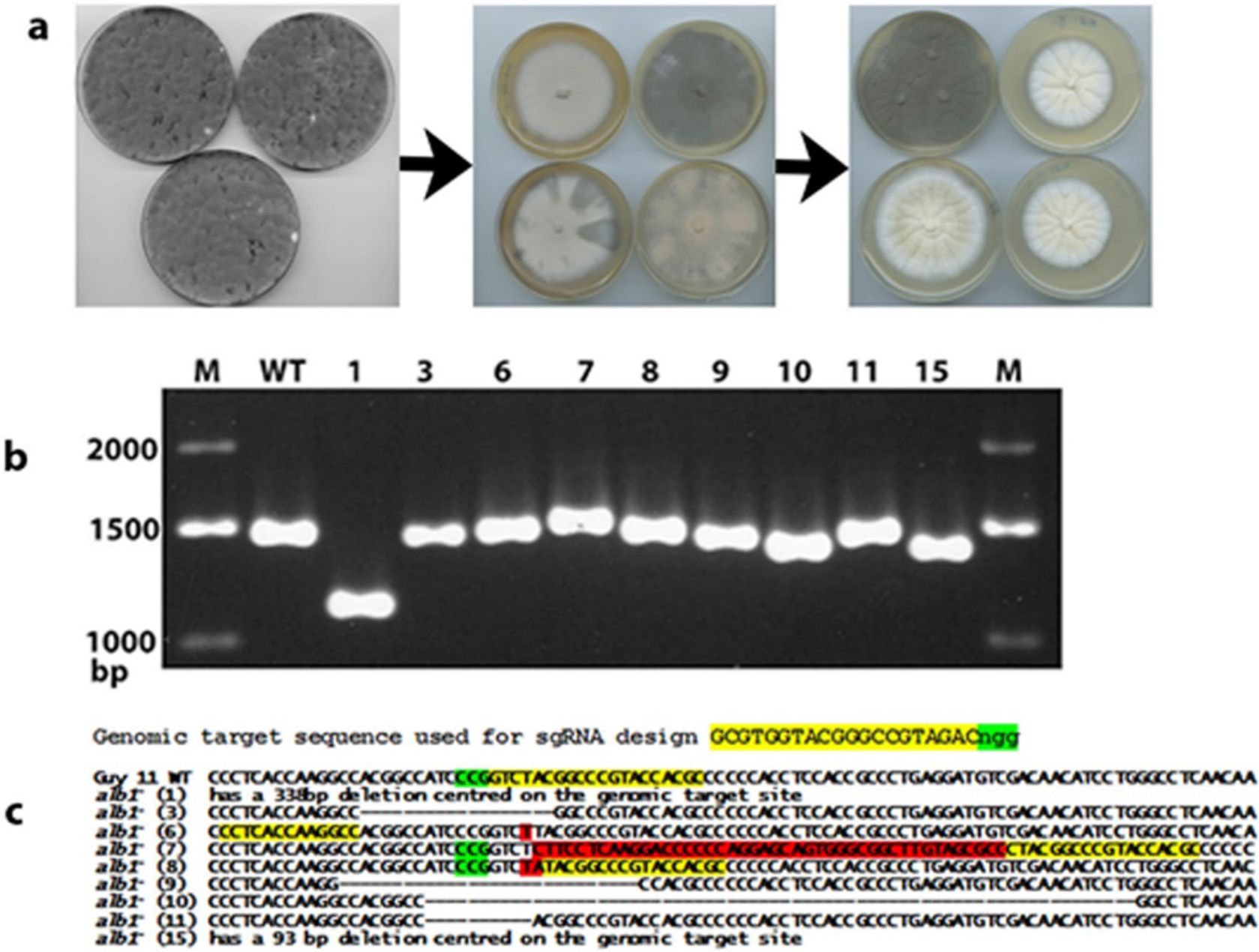
Marker-less RNP-CRISPR-Cas9 gene targeting of *ALB1* without donor DNA. **a.** Transformation plates from *ALB1*-targeting RNP-CRISPR-Cas9 without donor DNA showing rare albino patches and then followed by purification of albino regenerants by a combination of subculture (hyphal tip isolation) and single spore isolation to give pure albino colonies. **b.** Gel electrophoresis of the PCR products generated using the genomic DNA of the purified albino regenerants and primers PKS-ck-F and PKS-ck-R which flank the *ALB1*-targeting RNP-CRISPR-Cas9 genomic target sequence showing visible variation in product size. **c.** Sequences of the amplicons shown in **b**, showing a range of mutations and indels.

### Specific single nucleotide gene edits can be very efficiently accomplished using short oligonucleotide donor DNAs

One of the attractive features of CRISPR-induced gene editing is the ability to make highly specific changes to the coding sequence of a gene that could, for example, give rise to a single amino acid change in a protein product. To test whether single nucleotide edits were feasible using RNP-CRISPR-Cas9 in *M. oryzae*, we attempted to edit the gene *SDI1*, which encodes a subunit of the succinate dehydrogenase enzyme. We designed a *SDI1*-targeting RNP, to introduce a mutation that leads to an amino acid change in the enzyme known to confer resistance to the fungicide carboxin (see Fig. 3a; 22). At the same time we attempted to test how short homologous regions on the donor DNA can be, while still efficiently editing the target gene. By introduction of a *SDI1*-targeting RNP (Fig. 3b) and oligonucleotide donor DNAs of varied lengths (Fig 3a) into Guy 11 protoplasts, we were able to demonstrate that a 50-80 bp double stranded (ds) oligonucleotide donor DNA, containing the desired single base edit, was sufficient to efficiently edit *SDI1*, as shown in Fig 3. c and d. We therefore employed 80 bp donor DNAs in all of our subsequent experiments.

**Figure 3.**
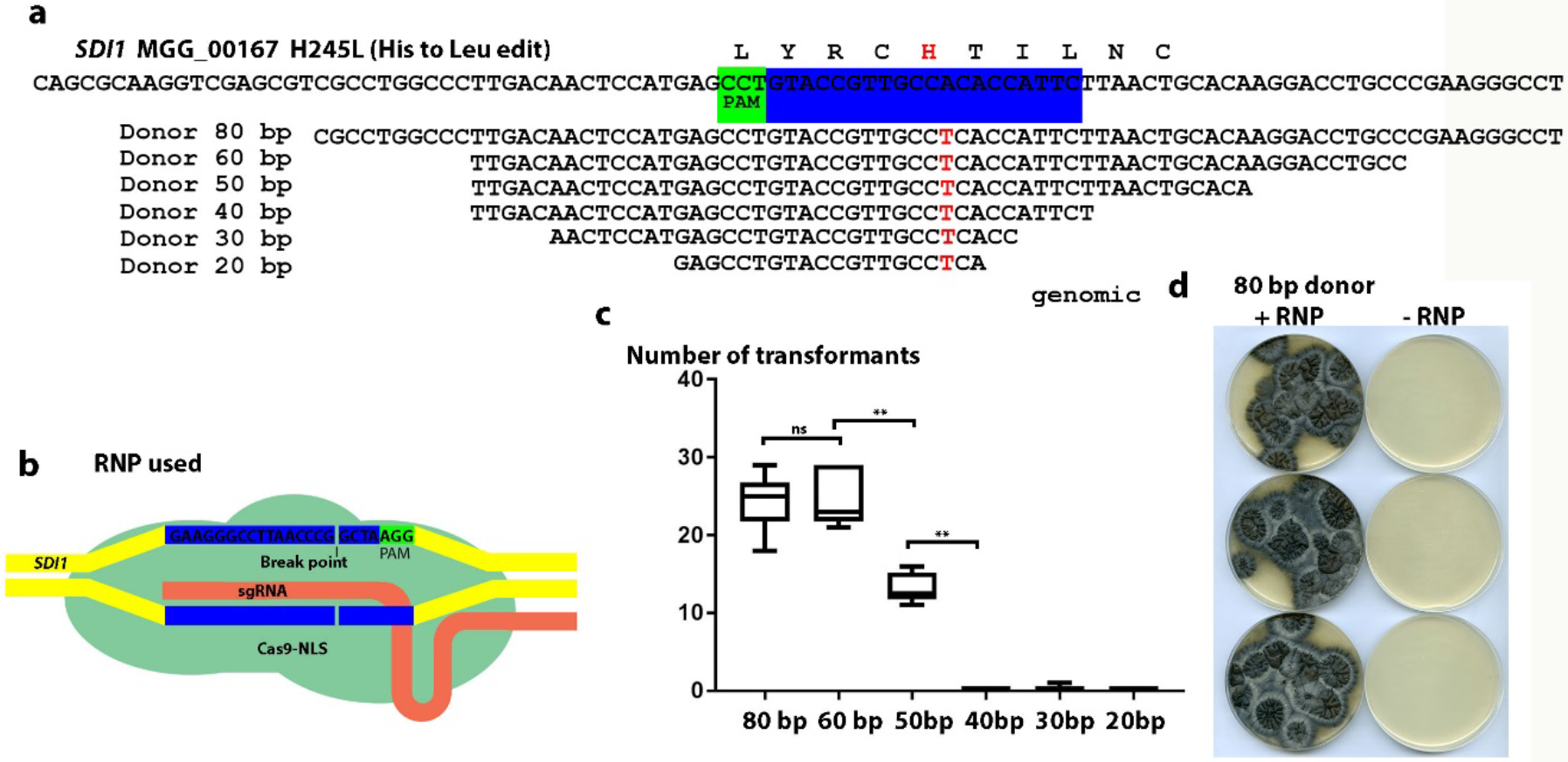
Oligonucleotide-mediated RNP-CRISPR-Cas9 gene editing of *SDI1* to confer carboxin resistance. **a.** Illustration of the substitution required to give a carboxin resistant form of the succinate dehydrogenase subunit product of MGG_00167 and the oligonucleotide donor DNAs capable of introducing the required one nucleotide change necessary tested. Also indicated is the genomic target sequence of the *SDI1*-targeting RNP-CRISPR-Cas9 complex employed. **b.** Diagram showing the RNP used and the predicted DSB at the *SDI1*-targeting RNP-CRISPR-Cas9 genomic target sequence. **b.** The number of transformants obtained using the donor DNAs in A in combination with the RNP illustrated in **b. d.** Transformants from the 80 bp long donor DNA shown in A. transformed together with the RNP complex illustrated in **c.** and also showing control plates where only the donor DNA without RNP was transformed (although no transformants are visible on the control plates, using the RNP + the 30 bp donor one carboxin resistant transformant was obtained which may indicate that very rarely the short oligos can recombine in the absence of the RNP complex; no other carboxin resistant transformants were obtained in the other controls).

### Development of a gene co-editing strategy in *M. oryzae*

We observed that mutants generated by NHEJ using RNPs without a selectable marker, did not arise at a frequency that would make gene targeting practicable. We therefore set out to develop a different method to enrich for gene-edited transformants. We decided to adopt a co-targeting approach in which two independent RNPs are transformed into *M. oryzae* together. We reasoned that a proportion of cells would take up both complexes and be edited at both loci. If one of the genes had an easily scorable phenotype that could be used as a selectable marker, we could therefore select transformants more easily and then determine whether the second locus had also been edited. As a proof of principle, we introduced RNPs targeting the succinate dehydrogenase subunit-encoding gene *SDI1* and *ALB1* simultaneously, together with the 80 bp oligonucleotide donor dsDNA, which we had already established was able to convert the *SDI1* gene to an allele bestowing carboxin resistance. We employed two different *ALB1-targeting* donor DNAs, both of which introduce a premature stop codon in the gene, close to its 5’ end (Fig. S1). One of the donor DNAs introduces the edit within the genomic target sequence, while the other is predicted to generate an edit 40 bp from the PAM, that would allow us to assess whether we can create edits at some distance from the DSB. We found that when Guy 11 was used as a recipient, these donor DNAs integrated in approximately 50% of the albino transformants, which represented 1-2 % of all the carboxin resistant transformants, as shown in Table S4. Furthermore, the donor DNA that introduces an edit outside the genomic target sequence was as efficient at editing as the other donor. The albino mutants that lacked the integrated donor DNAs exhibited indels, typically 3-4 bp 5’ from the PAM. These are indicative of mutations generated by NHEJ of the CRISPR-generated DSB. Surprisingly, in one instance an albino mutant arose by integration of the *SDI1*-targeting donor DNA at the *ALB1* locus. By contrast, when we employed a *Δku70* mutant (23) that lacks the NHEJ pathway, all albino mutants generated showed precise integration of the *ALB1* targeting donor DNA. Moreover, the efficiency of co-editing both loci increased, although the number of overall transformants was reduced (Table S4). To determine if this approach, which we henceforth refer to as co-editing, was applicable to other genes, we set out to co-edit both the *ILV2* and *TUB2* genes, which encode acetolactate synthase and b-tubulin, respectively. These genes can be edited at a single nucleotide to give rise to alleles encoding sulfonylurea and benomyl resistant mutants, respectively (see Fig. S2a; refs 24, 25;). RNPs were first created to introduce these edits and then tested individually (Fig. S2b). We then conducted a co-editing experiment by transforming the *ILV2* and *TUB2* targeting RNPs, together with the two corresponding oligonucleotide donors. We selected for sulfonylurea resistance and then calculated the proportion of sulfonylurea resistant transformants that were also benomyl resistant. Consistently, we observed ^~^1% efficiency of co-editing (Table S4). We were able to confirm the edits that had occurred in these transformants by direct sequencing of amplicons containing the target sequence. Mutations generated by NHEJ repair would in most conceivable instances, not be selected for by these experiments. Together, these experiments demonstrated that we are able to generate marker-less mutations in *M. oryzae* at target loci by employing a straightforward co-editing strategy.

### Co-editing allows generation of in-locus GFP-tagged gene fusions and conditional mutant alleles in *M. oryzae* without a selectable marker

To demonstrate that co-editing could be employed to generate novel genotypes in any gene of interest, we decided to tag the *SEP5* septin-encoding gene (26) with GFP, using CRISPR co-editing at the native locus. We also tested whether we could exploit co-editing to introduce a two nucleotide edit into the *SEP6* septin-encoding gene to create a temperature-sensitive allele. We generated a G234E substitution into *SEP6*, which corresponds to a mutation (G247E) that in the Sep6 orthologue Cdc12 in *Saccharomyces cerevisiae*, gives rise to a temperature sensitive (ts) form of the septin (Fig. 4a; 27). Using CRISPR-mediated co-editing we were able to generate both a *sep6*^*G234D*^ allele (Fig. 4b and c) and a *SEP5-GFP* strain, as shown in Fig. 4D and E. To confirm that the correct genotype had been created at the corresponding loci, we sequenced amplicons of both genomic regions (Fig. 4b). The *SEP5-GFP* strain was identified by examination of 200 transformants, while the *sep6*^*G234D*^ mutation was identified among 79 transformants. These experiments confirmed that co-editing can be employed to rapidly and precisely manipulate genes in *M. oryzae*. We confirmed that the *sep6*^*G234D*^ mutation leads to a temperature sensitive loss of virulence, as shown in Fig. 4c. The *sep6*^*G234D*^ mutant was unable to cause rice blast symptoms at the non-permissive temperature of 30°C (Fig. 4c). Replacement of *SEP5* with a *SEP5-GFP* gene fusion at the native locus meanwhile leads to visualisation of a GFP-tagged septin ring at the *M. oryzae* appressorium pore (Fig. 4e), identical to that previously reported for an ectopically integrated gene fusion (26).

**Figure 4.**
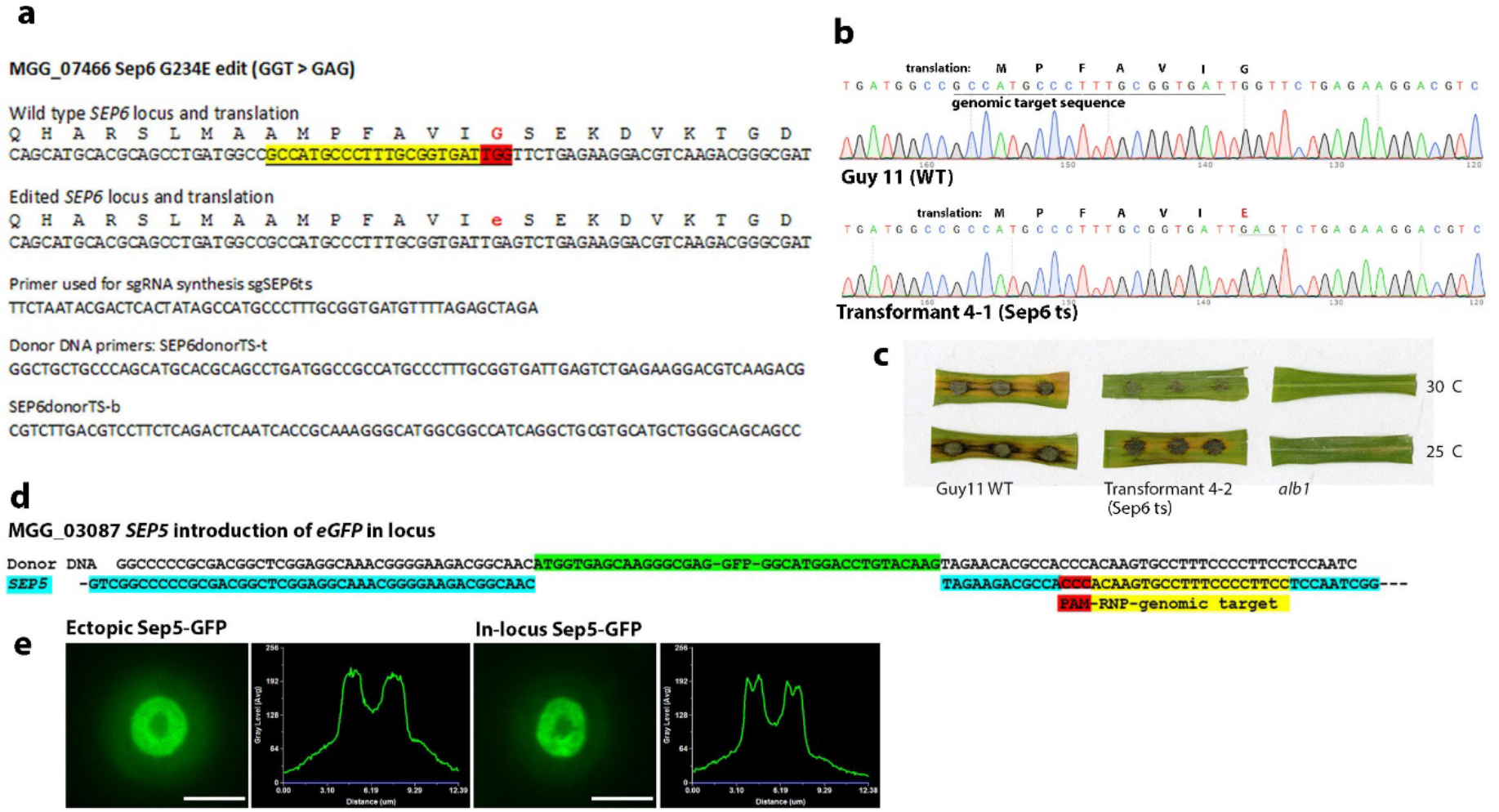
RNP-CRISPR-Cas9 gene co-editing to introduce a N-terminal GFP tag to Sep5 and a temperature sensitive mutation in Sep6. **a**. An illustration of the genomic target sequence for generation of a Sep6 temperature sensitive (ts)-encoding allele of *SEP6* by co-editing at the native locus. **b**. An illustration of the genomic target sequence for generation of a strain where the GFP-encoding gene is inserted after the START codon of the Sep5-encoding gene potentially giving rise to a *SEP5-GFP* expressing strain with an in locus replacement of the native *SEP5* gene. **c.** Confirmation of the introduction of the desired mutation using sequencing of a 472 bp amplicon generated using the primers SEP6ts?-f CACACCCTGAAGCCCCTTGATATC and SEP6ts?-R CTCCTCGGTTGTGTGGATGAG. **d**. Leaf sections showing the infection of rice cultivar Co-39 with conidia of a strain with the ts allele of *SEP6*. **e**. Micrographs showing the appressorial Sep5-gfp containing septin ring in a strain where *SEP5* was replaced by the Sep5-gfp-encoding gene in locus by RNP-CRISPR-Cas9 co-editing and the corresponding septin ring in the ectopically integrated *SEP5-GFP*-expressing strain constructed by Dagdas and co-workers (26). The linescan graphs show the Sep5-GFP fluorescence in a transverse section of the individual appressoria shown in the micrographs.

### A novel selection strategy allowing construction of isogenic gene edited mutants in *M. oryzae*

Gene manipulation in *M. oryzae*, as in all plant pathogenic fungal species, has normally involved the generation of mutants that also express selectable marker-encoding genes. The effect of expression of antibiotic resistance genes may be negligible in most instances, but is still likely to have consequences, which for the most part remain unknown. Ultimately, it would be desirable to generate a mutant that contains a specific edit, but that is in all other respects isogenic to the progenitor strain. We reasoned that the reported negative cross resistance of certain benomyl resistant mutations to the compound diethofencarb (28, 29), might provide a novel counterselection strategy that would allow us to generate isogenic CRISPR mutants in *M. oryzae*. We were able to confirm in plate growth tests that the wild type *M. oryzae* strain, Guy11, can exhibit normal growth in the presence of 10 μg mL^-1^ diethofencarb, whereas a benomyl-resistant strain harbouring a *TUB2* allele with a E198A mutation cannot grow under the same conditions (Fig. 5A). We therefore used a TUB2-targeting RNP and an 80 bp oligo donor that restores the *TUB2* sequence to wild type, and introduced this into a benomyl resistant transformant, previously created using RNP-CRISPR-Cas9. This led to the generation of diethofencarb-resistant transformants at a very high frequency (Fig. 5c). The diethofencarb-resistant transformants were as sensitive to benomyl as Guy 11. No diethofencarb-resistant transformants were generated in the absence of the TUB2-targeting RNP. Furthermore, we observed that the counterselection is very tight, because no wild type *TUB2* strains can grow on 10 μg mL^-1^ benomyl, and no benomyl-resistant *TUB2*^*El98A*^ strains are able to grow at all on 10 μg mL^-1^ diethofencarb as shown in Fig. 5a. We conclude that CRISPR-Cas9-RNP mediated generation of *TUB2*^*E198A*^ benomyl resistant strains of *M. oryzae*, followed by a second round of CRISPR-Cas9-RNP to restore a wild type *TUB2* sequence, bestowing diethofencarb resistance, provides a means of generating isogenic, markerless, gene-edited mutants. The counter selection strategy also represents an excellent way by which to build multiple mutations in a single strain of the fungus, as it is straightforward to cycle between the two states, and thereby introduce further specific edits or other manipulations each time by co-editing.

**Figure 5.**
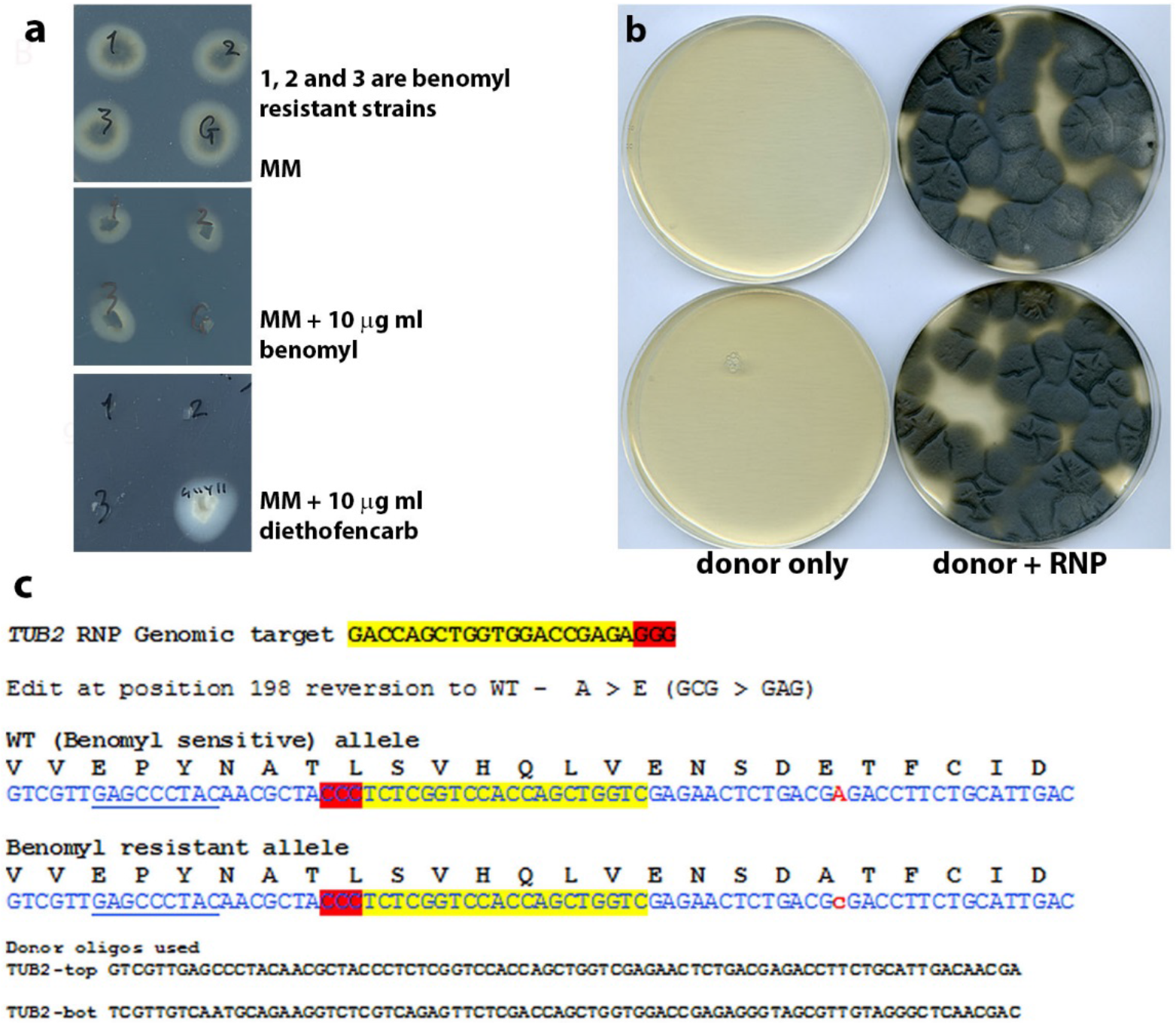
Counterselection exploiting the diethofencarb sensitivity of strains expressing E198A beta-tubulin. **a**. Plates illustrating the negative cross resistance of benomyl resistant transformants to diethofencarb. 1, 2 and 3 are three different benomyl resistant strains generated using RNP-CRISPR while G (Guy 11 is the benomyl sensitive control). **b.** Plates showing the transformation of the benomyl resistant strain number 2 with the *TUB2*-targeting RNP-CRISPR-Cas9 and a donor DNA (shown in C) conferring the reversion to the wild type *TUB2* sequence and diethofencarb resistance. **c**. Illustration of the genomic target sequence for the RNP used and the sequence edit for reversion to a WT TUB2 sequence using the donor DNA shown.

### Assessment of off target effects in albino mutants generated by RNP-CRISPR without donor DNA

One potential constraint on the use of CRISPR-Cas9 gene editing is the possibility of off-target mutations at sites showing significant similarity to the genomic target sequence (30). Although no mismatch is tolerated in the seed sequence proximal to the PAM, in some species up the 5 bp mismatches are tolerated at the PAM-distal end of the genomic target sequence, resulting in potential for off-target cleavage (31). For this reason, most programmes used for automated protospacer selection search for a protospacer with minimal potential for off target cleavage. The degree of off target cleavage, however, varies considerably between organisms and is, of course, also a function of genome size. However, careful sgRNA design and limited longevity of the Cas9-SgRNA complex in a cell, seem to be major means to maintain editing specificity (31). An advantage of the using RNP-CRISPR gene editing is that there is evidence that the transient presence of the CRISPR machinery in the cell may also limit off target effects (32).

In order to assess whether off target mutations had occurred in CRISPR generated mutants, we randomly selected two *alb1* mutants generated by CRISPR-Cas9-RNP and, as a control, two strains that had been through the transformation procedure and regenerated but not exposed to RNP complexes. The genomes of these *M. oryzae* strains were sequenced (Table S5). The presence of SNPs and small INDELs was then determined in *alb1* mutant strains, compared to the control strains. Potential off-site mutations were detected based on the presence of an insertion or deletion (INDEL) within 3bp of the PAM site or in regions showing sequence identity at the mutant position, of at least 10 bases to the guide RNA. We found that the *alb1*_3 mutant had 44 SNPs and 30 INDELS, while the *alb1*_6 mutant had 40 SNPs + 33 INDELS across the whole genome, when compared to the control strains, which represent the sequence of the progenitor Guy11 isolate. Our analysis showed that there are 115 sites in the *M. oryzae* genome with at least 10 bp identity to the gRNA, but we found that none of these corresponded to mutations detected in either of the *alb1* mutants. We conclude that no off-target CRISPR generated mutations occurred in the *alb1* mutants of *M. oryzae*.

## Discussion

In this report, we have demonstrated the efficient generation of CRISPR induced gene edits in the rice blast fungus using purified CRISPR machinery components. We were motivated to develop this method because stable expression of Cas9 appears to be very toxic to *M. oryzae* in the same way as reported in some other species, including fission yeast (33, 34, 35). The transient nature of CRISPR-Cas9-ribonucleoprotein expression provides an excellent means of circumventing the problems associated with Cas9 toxicity and rapidly generating gene edited mutants in *M. oryzae*. This contrasts with the previously published method requiring expression of the Cas9-encoding gene (14), which we were unable to reproduce in spite of extensive efforts. The key advantages of RNP-mediated CRISPR-Cas9 editing are its efficiency, accuracy, and especially its speed. Use of oligonucleotide-based or PCR-amplified donor DNAs obviates the need for labour-intensive DNA cloning and thereby dramatically reduces the time and cost required to make precise gene manipulations in this species. Additionally, we demonstrated that RNP-CRISPR editing is highly specific, because we saw no evidence of off-target mutations in the genomes of two CRISPR generated mutants. RNP-CRISPR therefore is an extremely useful and adaptable addition to the *Magnaporthe* gene manipulation toolbox as a simple amendment to existing transformation protocols. Furthermore, as the price of the necessary components of the RNP complex falls over time, we predict that RNP-CRISPR-Cas9 editing will become a standard manipulation in a very short time in *M. oryzae*.

We observed that although mutations resulting from RNP-CRISPR and NHEJ-dependent repair were possible without a selectable marker, these occurred at a frequency too low to be practically exploited. One interpretation of this result is that NHEJ may be highly accurate in *M. oryzae*, but a more likely explanation is that a very small proportion of fungal protoplasts actually take up the RNP. It is therefore possible that RNP complexes might be more efficiently delivered by other means, such as electroporation or biolistic delivery. We were, however, able to overcome this potential limitation by developing a gene co-targeting strategy, that we termed co-editing. This co-editing approach significantly enriches for specific edits in a marker-less fashion−without introduction of a further selectable marker sequence at the locus of interest. Co-editing was found to occur in a useful number (around 1%) of transformants that were also edited to bestow carboxin, benomyl or sulfonylurea resistance, respectively. This required a second RNP and donor DNA targeting a gene of interest, in addition to the RNP donor pair generating the mutation to confer antibiotic resistance. Using oligonucleotide donor DNAs also had the additional advantage that it generated very large numbers of antibiotic-resistant transformants, from which it was straightforward to select co-edited mutants.

When the gene co-editing method is combined with the benomyl/diethofencarb-based counterselection strategy, we have provided a mechanism to generate gene edited mutants in *M. oryzae* that are truly isogenic to a progenitor wild type. This is very advantageous, not only for basic research (where studying a single mutation in isolation from any other genome pertubations is the best possible method), but also in fungal biotechnological applications where the lack of a resistance gene marker is important from a regulatory perspective. Although the efficiency of gene co-editing that we report here is rather low, it is likely that optimisation by adjustment of the ratio of the RNPs and/or donor DNAs may be possible in future. Furthermore, in the current report we show that 1 or 2 base changes are possible with an 80 bp donor DNA, but preliminary results in our laboratory indicate that similarly sized donor DNAs can efficiently and precisely delete small sections of genes of around 50 bp, that would facilitate simple PCR-based screens for gene-inactivated mutants. Additionally, the generation of small deletions may be a more attractive method for high-throughput gene functional analysis because they may be more stable than changes to a single nucleotide. It may also be possible to devise a screening strategy for co-edited mutant, based on PCR at very stringent conditions, or by coupling PCR with restriction digestion. However, the time saved by not having to construct vectors for gene manipulation makes co-editing an attractive option, even if it necessitates sequencing 100 or more transformants to identify the specific mutant required.

Our study does, however, raise some important questions too, that we will address in future. It is apparent, for instance, that some RNPs work more efficiently than others as has been reported in other species (36, 37). In making the most efficient use of co-editing, it is important that we better understand which protospacers are likely to generate the best results. Additionally, it has been suggested that protoplast-mediated transformation is in itself, mildly mutagenic (38), and in the long term it would be worth investing time to explore other means to deliver RNP complexes to *M. oryzae*. Electroporation, for example, might represent a less disruptive means of delivering the RNP-CRIPSR-Cas9 complex. If RNP complexes can be delivered at the same time as DNA donors via electroporation, it would, for instance, be possible to directly compare SNPs and indel frequencies in the genomes of fungal strains manipulated by these different methods to test whether electroporation or other delivery systems really are less mutagenic than protoplast generation.

In summary, the current study demonstrates that RNP-CRISPR-Cas9 gene editing is a simple, precise, reproducible, and rapid means by which gene manipulation can be carried out in *M. oryzae*. It is our hope that researchers investigating *Magnaporthe* species and related fungi will adopt the procedures described here and that this will empower researchers and accelerate discovery towards understanding and ultimately combatting the devastating rice blast fungus *M. oryzae* and similarly important pathogenic fungal species.

## Materials and methods

### Strains and Culture conditions and infection assays

The wild type strain Guy 11 and the NHEJ deficient mutant *Δku70* were grown in a controlled temperature room at 25°C with a 12h light/dark cycle. For tests of temperature sensitivity an incubator at 30°C with a 12h light/dark cycle was used to conduct cut leaf infection assays. Infection assays used two cm long leaf sections cut from 3 week old leaves of the dwarf indica rice cultivar CO-39 and were assessed after 5 days incubation at either 25°C or 30°C with a 12h light/dark cycle. For glufosinate and sulfonylurea selection, we used BDCM medium (39). For standard growth, we used CM (40). For selection using hygromycin, benomyl, carboxin or diethofencarb we used OCM which is CM osmotically stabilised with 0.8M Sucrose. Selective agents were used at a final concentration of 200 μg mL^-1^ for hygromycin or 150 μg mL^-1^ for sulfonylurea (chlorimuronethyl) 40 μg mL^-1^ glufosinate or 10 μg mL^-1^ benomyl 50 μg mL^-1^ carboxin or 10 μg mL^-1^ diethofencarb.

### SgRNA synthesis and RNP formation

SgRNA for complexing with Cas9-NLS was synthesised using the EnGen sgRNA synthesis kit available from New England Biolabs (NEB #E3322), according to the instructions provided, and purified before complexing to Cas9 using the RNA Clean & Concentrator-25 kit (Zymo Research). Cas9-NLS was purchased from New England Biolabs (NEB Catalog #: M0646M). Cas9-NLS was complexed with the sgRNA by a 10 minute incubation at room temperature. Oligonucleotides for use as templates in the reaction were purchased desalted from Eurofins Genomics UK. The sgRNA selection was carried out using the E-CRISP program at http://www.e-crisp.org/E-CRISP/ (41).

### Fungal transformation and introduction of RNPs and donor DNAs

PEG-mediated fungal transformation of Glucanex-generated protoplasts was achieved, as previously described (40). The RNP complexes and donor DNAs were introduced together before the step in the standard transformation procedure, where 50% PEG is added and the mixture then incubated for 25 min. During optimisation for reproducibility of the method, sgRNAs were always freshly synthesised and purified on the day of transformation and kept on ice following formation of the RNP complex. All experiments except co-targeting experiments using the SDI1-targeting RNP, used 2 μg of donor DNA and 6 μg of Cas9 mixed at a 1:1 molar ratio with the respective sgRNA (pre-complexed together for 10 min at RT) and protoplasts, in a volume of 150 μl at a concentration of 1.5 × 10^8^ protoplasts/ml. For the co-targeting experiments using the SDI1-targeting RNP, we used 1 μg of Cas9 precomplexed with 0.2 μg of sgRNA. For *Agrobacterium*-mediated transformation of Cas9-containing vectors, conidia of Guy11 were transformed as previously described (42).

### Cloning of *ALB1* and *RSY1* targeting donor DNAs

Oligonucleotides used in this study are listed in Table S6. To generate cloned donor DNA for repair of RNP-CRISPR generated DSBs, in the case of *ALB1* we amplified a 1.4 kb *ALB1* fragment using the primers ALB1-for-EcoRI and ALB1-rev-SpeI. The resultant PCR product was digested with EcoRI and XhoI gel purified and ligated to EcoRI + SalI digested pCAMBIA0380 thus destroying the SalI site in this vector multi-cloning site. The resultant vector pCAMB-ALB1 was linearized using a I site in the middle of the *ALB1* fragment and ligated to the *HPH* cassette, conferring hygromycin resistance excised from plasmid pCB1636 (39) using SalI to create pCAMB-ALB-HPT. A donor DNA was generated by PCR amplification of the *HPH* interrupted *ALB1* fragment from pCAMB-ALB-HPT using the primers ALB1-for and ALB1-rev. In the case of *RSY1*, RSY-donor-f and RSY-donor-r were used to amplify a 1.8 kb *RSY1* fragment which was cloned into the vector pGEMT-easy (Promega) to give pGEM-RSY. pGEM-RSY was digested with XhoI and ligated to a Sa1I *HPH* fragment to give pGEM-RSY-HPT which could then be used as a template for amplification of the required donor DNA (*HPT* interrupted *RSY1*) using RSY-donor-f and RSY-donor-r.

### Cloning of *Agrobacterium tumefaciens* compatible Cas9 expressing vectors

To generate an *Agrobacterium* compatible vector to introduce Cas9 and an *ALB1* targeting sgRNA transcribing sequence into *M. oryzae*, we used the nuclear localised, Cas9 codon-optimised for *N. crassa* in vector p415-PtrpC-Cas9-TtrpC-CYC1t (43) as a template for high fidelity PCR amplification of Cas9-NLS under control of the *Aspergillus nidulans TrpC* promoter and terminator sequences, using the primers Cas9-recom-f and Cas9-recom-r. We then used yeast recombination in *Saccharomyces cerevisiae* strain DS94 (*MATα*, *ura3*-52, *trp1*-1, *leu2*-3, *his3*-111, and *lys2*-801 (44)) to recombine this fragment into a 12,869 bp XhoI-BamHI fragment of the “soft-landing” vector pS315 pMMag_Cbx-mCherry, which will integrate at the *SDI1* locus as a single copy to confer carboxin resistance (pS315-pMMag-Cbx-mCherry was a kind gift from Prof. Gero Steinberg, University of Exeter). The resultant vector p315-Cas9-csr-1 was then further modified to introduce sgRNA transcribing sequences targeting *ALB1* or *RSY1* This was achieved by generating a PCR-amplified fragment containing the desired guide RNA encoding sequence created by using the plasmid p426-SNR52p-gRNA.csr-1.Y-SUP4t as a template, with primers PKS1gRNA-f and PKS1gRNA-r for *ALB1* and in the case of *SDH1* the PKS1gRNA-r primer was replaced with the primers SDH1-sg-r. This PCR results in the replacement of the *Neurospora crassa csr-1* targeting guide with the *M. oryzae ALB1* or *RSY1* targeting guide under the control of the *SNR52* promoter. The resultant amplicons were then used as a template to generate recombination competent frgaments using the primers gRNAto315Cas-R and sgRNA-rec-R. The products of this amplification were recombined separately in yeast into XbaI digested p315-Cas9-csr-1. Correct assembly of the vectors in yeast was confirmed following extraction from yeast strains, transformation of, and purification from *E. coli* followed by analysis by restriction digests and partial sequencing. The vectors p415-PtrpC-Cas9-TtrpC-CYC1t and p426-SNR52p-gRNA.csr-1.Y-SUP4t were gifts from Christian Hong (Addgene plasmid numbers 68059 and 68060 respectively).

### Bioinformatic analysis for detection of potential off target mutations

Genomic DNA was sequenced on Illumina HiSeq 2500 using standard reagents and protocols producing 125 base paired-end reads. Reads were filtered using the fastq-mcf program from the ea-utils package, eautils (45): “Command-line tools for processing biological sequencing data”; https://github.com/ExpressionAnalysis/ea-utils). Genomes were assembled ‘de novo’ using SPAdes 3.11.0 (46). For SNP and INDEL calling reads were aligned against the *M. oryzae* reference genome (47) using BWA (48). SNPs and INDEL were identified with a bespoke pipeline using bcftools and vcfutils from the SAMtools package as well as SnpSift (49, 50). SNPs called from the two *alb1* mutants were compared to two regenerated strains that had been subjected to the transformation protocol but not exposed to RNPs, based on analysis made by Schuster and co-workers (38).

## Data availability

All data generated or analysed during this study are included in this published article (and its Supplementary Information files) or are available from the corresponding author on reasonable request.

## Acknowledgements

This work was funded by a European Research Council Advanced Investigator Award to NJT under the European Union’s Seventh Framework Programme (*FP7/2007-2013*)/ ERC grant agreement n° 294702 GENBLAST and by a BBSRC grant (BB/BB/N009959/1) to NJT.

**Supplementary information** accompanies this paper

**The authors declare no competing interests.**

## Author Contributions

N.J.T. and A.J.F. designed the experiments, oversaw the study, and wrote the manuscript. A.J.F., M.M.U., X.Y., and S.W. carried out experimental work. D.S. conducted the genomic analysis of off-target effects.

